# Scalable, methanol-free manufacturing of the SARS-CoV-2 receptor binding domain in engineered *Komagataella phaffii*

**DOI:** 10.1101/2021.04.15.440035

**Authors:** Neil C. Dalvie, Andrew M. Biedermann, Sergio A. Rodriguez-Aponte, Christopher A. Naranjo, Harish D. Rao, Meghraj P. Rajurkar, Rakesh R. Lothe, Umesh S. Shaligram, Ryan S. Johnston, Laura E. Crowell, Seraphin Castelino, Mary Kate Tracey, Charles A. Whittaker, J. Christopher Love

## Abstract

Prevention of COVID-19 on a global scale will require the continued development of high-volume, low-cost platforms for the manufacturing of vaccines to supply on-going demand. Vaccine candidates based on recombinant protein subunits remain important because they can be manufactured at low costs in existing large-scale production facilities that use microbial hosts like *Komagataella phaffii* (*Pichia pastoris*). Here, we report an improved and scalable manufacturing approach for the SARS-CoV-2 spike protein receptor binding domain (RBD); this protein is a key antigen for several reported vaccine candidates. We genetically engineered a manufacturing strain of *K. phaffii* to obviate the requirement for methanol-induction of the recombinant gene. Methanol-free production improved the secreted titer of the RBD protein by >5x by alleviating protein folding stress. Removal of methanol from the production process enabled scale up to a 1,200 L pre-existing production facility. This engineered strain is now used to produce an RBD-based vaccine antigen that is currently in clinical trials and could be used to produce other variants of RBD as needed for future vaccines.

## Manuscript

As new variants of SARS-CoV-2 emerge, continued development of diagnostics, vaccines, and reagents remains essential to address the COVID-19 pandemic. The SARS-CoV-2 spike protein is an essential reagent for serological assays, and a component of several proteinbased vaccines (Guebre-Xabier et al., 2020; Tian et al., 2020). Vaccine candidates based on protein subunits are also important ones for enabling interventions for the pandemic in low- and middle-income countries (LMICs) due to existing large-scale manufacturing facilities and less stringent temperature and storage requirements for distribution (Dai et al., 2020). We and others have reported vaccine designs based on the receptor binding domain (RBD) of the spike protein (Dalvie et al., 2021). In these designs, the RBD can be produced independently, and subsequently displayed on protein or lipid nanoparticles for enhanced immunogenicity (Cohen et al., 2021; Walls et al., 2020). The 201 amino acid RBD is an especially promising antigen for accessible vaccines because it can be manufactured at low cost and high volumes in microbial hosts (Chen et al., 2020; Pollet et al., 2020). Here, we report an engineered yeast strain with enhanced secretion of the SARS-CoV-2 RBD from the circulating variants of Wuhan Hu-1, B.1.1.7, and B.1.351 strains of the virus. This engineered host has been successfully deployed at 1,200 L scale to produce a vaccine component currently in clinical trials.

The methylotrophic yeast *Komagataella phaffii* (*Pichia pastoris*) is routinely used for the production of therapeutic proteins at large volumes because of its high-capacity eukaryotic secretory pathway (Love et al., 2018). Another key advantage of this production host is the strong, tightly regulated, methanol-inducible promoter, P_AOX1_, used for expression of the recombinant gene (Ahmad et al., 2014). This promoter enables outgrowth to high cell densities with inexpensive feedstock like glycerol before induction of the recombinant gene with methanol feed. Methanol can pose challenges, however, in large-scale facilities, including high heat generation during fermentation and flammability concerns while in storage (Potvin et al., 2012). The impact of these challenges is that facilities require specific designs or modifications to handle methanol. This requirement could limit the number of manufacturing facilities available for the production vaccine components like the RBD antigens in *K. phaffii* in a pandemic. We sought to reduce or eliminate the requirement for methanol for efficient secretion of the RBD.

We previously reported the production of the SARS-CoV-2 RBD (Wuhan-Hu-1 sequence) in an engineered variant of *K. phaffii* (Brady et al., 2020; Dalvie et al., 2021). To assess the feasibility of methanol-free production, we cultivated the strain expressing RBD regulated under the native AOX1 promoter, and induced expression of the recombinant gene with varying amounts of methanol (Fig. 1A). Interestingly, the secreted titers of RBD increased as the concentrations of methanol were reduced. We also induced protein production with a combination of methanol and sorbitol—a supplementary carbon source that does not repress P_AOX1_ expression—and observed a further increase in titer.

**Fig. 1.**
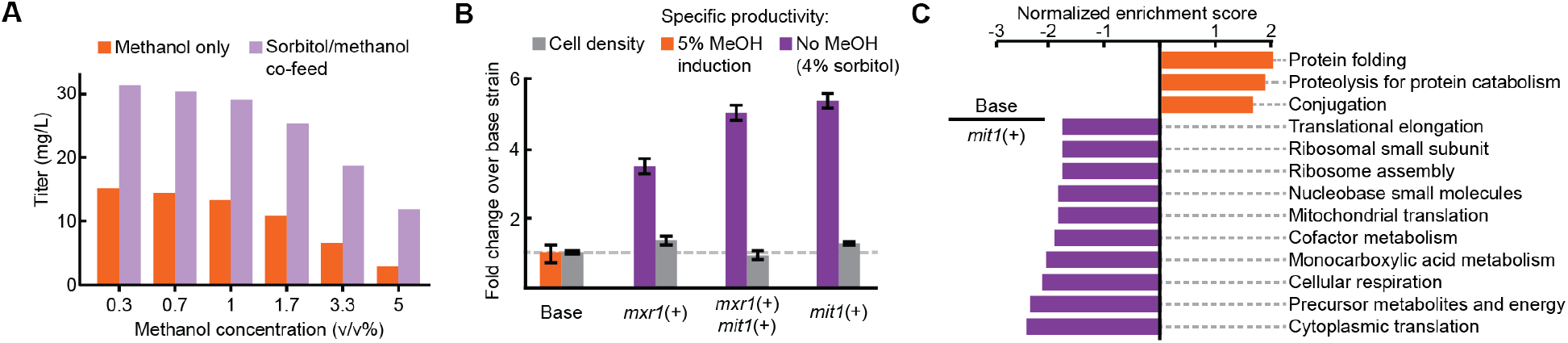
Improved productivity and decreased stress in methanol-free RBD expression (A) Titer of RBD secretion from the base strain in 3 mL plate culture. (B) Performance of three engineered strains in 3 mL plate culture. (C) Enriched gene sets between the base strain (orange) and the *mit1*+ strain (purple).

Given these results with reduced quantities of methanol in these batch cultivations, we hypothesized that we could achieve efficient secretion of the RBD with no methanol. Expression of genes regulated by P_AOX1_ in wild type *K. phaffii* in the absence of methanol is inconsistent, even with non-repressive carbon sources like sorbitol (Vogl et al., 2018). Several studies, however, have demonstrated that constitutive overexpression of activating transcription factors can lead to consistent activation of P_AOX1_ without methanol (Shi et al., 2019; Vogl et al., 2018). To test production of RBD without methanol, we integrated additional copies of the endogenous transcription factors *mit1* and *mxr1* into the *K. phaffii* genome under a glycerol-repressible promoter (Dalvie et al., 2019). We cultivated these strains for protein production by feeding with only sorbitol (Fig. 1B). We observed a >3-fold increase in specific productivity in all strains, particularly with a strain containing only one extra copy of the transcription factor *mit1* (>5-fold). To assess the potential source of improved productivity, we examined the transcriptomes of the methanol-fed initial strain and the modified, sorbitol-fed *mit1*+ strain by RNA-sequencing, and analyzed the variations by gene set enrichment analysis (Fig. 1C). We observed significantly higher expression of genes associated with protein folding stress in the methanol-fed condition compared to the sorbitol-fed *mit1*+ condition (family-wise error p=0.003). These results suggested that sorbitol-fed *mit1*+ may improve productivity by mitigating protein folding stress associated with RBD production.

After comparing the specific productivity of the methanol-free strain (*mit1*+) to the methanol-induced (base) strain, we assessed the production of RBD using both strains on InSCyT, a continuous, automated, perfusion-based manufacturing platform (Crowell et al., 2018). The base strain exhibited low titers (~30 mg/L) in perfusates and significant cell lysis after ~120 h of fermentation in perfusion (Fig. 2A-B). In contrast, the *mit1*+ strain maintained protein secretion at >50 mg/L/day for the duration of a >200 h campaign. RBD purified from the perfusates produced by the base strain also contained more host-related impurities than RBD from the *mit1*+ campaign (Fig. 2C). These results from the sustained production of RBD, including the cell lysis observed in the base strain, are consistent with the observations for increased cellular stress relative to the *mit1*+ strain, and suggest the transcriptional changes observed also translated into variation in protein expression as well.

**Fig. 2.**
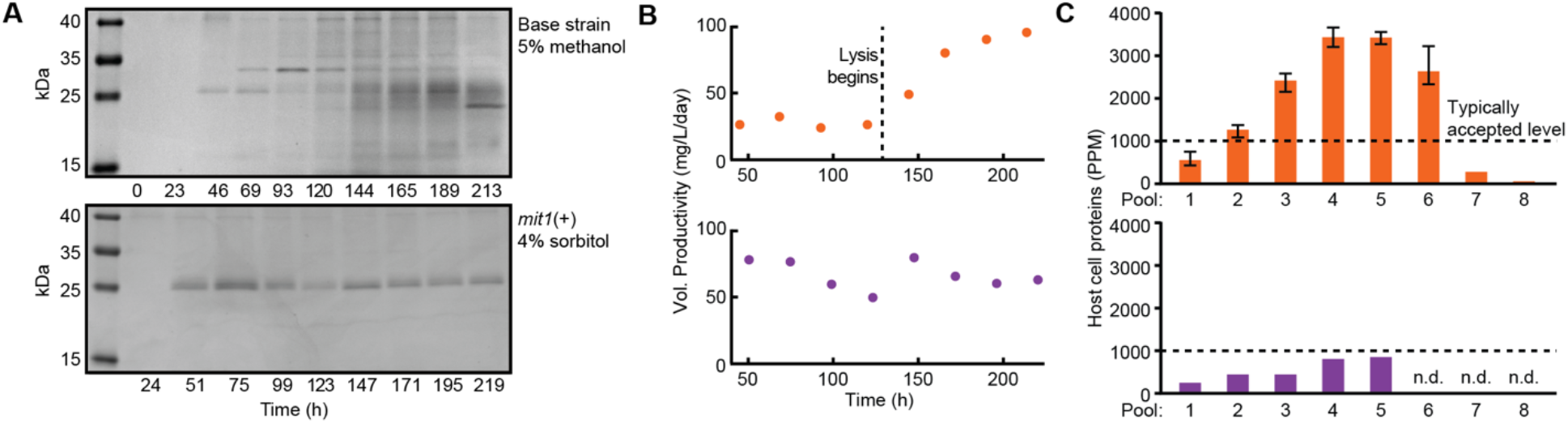
Sustained productivity of the methanol-free strain in perfusion fermentation (A) Reduced SDS-PAGE of upstream reactor samples for the duration of each campaign. (B) Upstream reactor titer of RBD. (C) Host cell protein concentrations in purified pools of RBD, measured by ELISA.

From these data for the improved production of RBD in bioreactors with the modified strain without methanol, we then generated an *mit1*+ strain that expressed RBD with a C-terminal fusion of SpyTag, a short peptide that can mediate a transpeptidation reaction with a cognate SpyCatcher polypeptide, which can be presented on protein nanoparticles for example (Reddington and Howarth, 2015). We expressed and purified the RBD-Spytag from this strain in a 200 mL shake flask culture. We also transferred this *mit1*+ strain encoding RBD-SpyTag, to a facility for GMP manufacturing in a 1,200 L fed batch process. In this process, the strain produced 21 mg of purified, clinical quality RBD-SpyTag per liter of fermentation, or approximately >1 million doses from a single reactor batch, assuming a vaccine formulation with 25 μg of RBD-SpyTag per dose. The two purified products were similar by SDS-PAGE, and exhibit nearly identical glycan profiles, indicating consistency in the quality attributes of the molecules produced at these two scales with this modified strain for methanol-free production (Fig. 3).

**Fig. 3.**
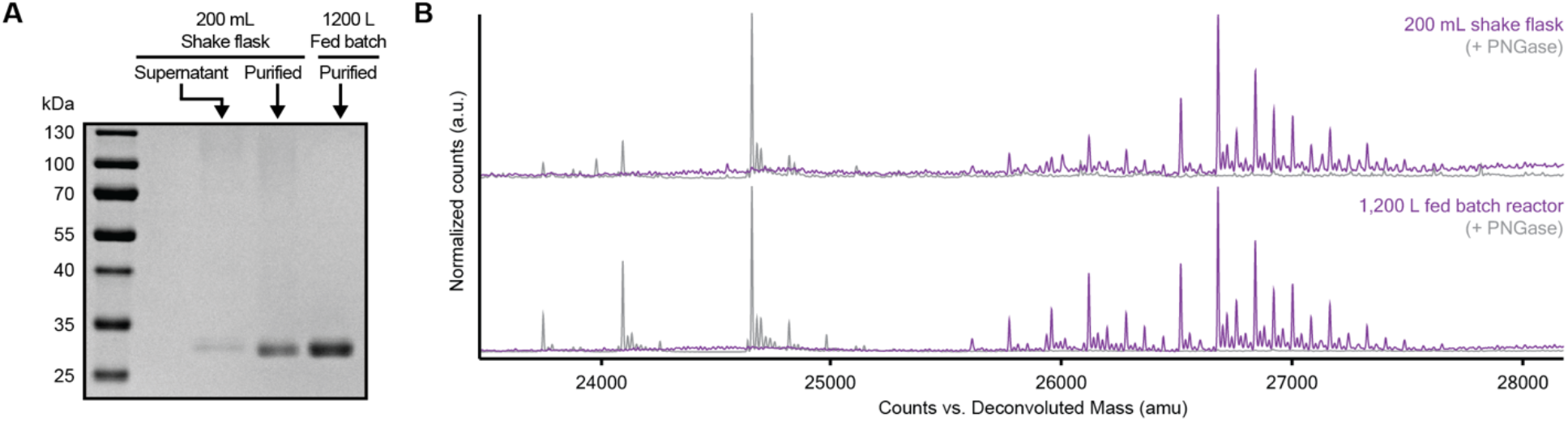
RBD-SpyTag produced at lab scale and GMP scale A) Reduced SDS-PAGE of RBD-SpyTag in crude shake flask supernatant, purified from shake flask cultivation, and purified from a fed batch process. B) Intact mass spectra of purified RBD-SpyTag from each manufacturing process. Overlayed spectra are before and after treatment with PNGase.

We next sought to assess whether or not this modified *mit1*+ strain could improve the production of sequence variants for other circulating SARS-CoV-2 virus strains as well. We generated strains expressing RBD-B.1.1.7 and RBD-B.1.351 in both the base and *mit1*+ strain backgrounds, and evaluated their specific productivities in different media for production (Fig. 4). In all strains, reduced methanol feed improved productivity. For all RBD variants, only *mit1*+ engineered strains maintained improved productivity in the absence of methanol. This result demonstrates that the engineered *mit1*+ strain could facilitate new cell lines for manufacturing other RBD variants without methanol for seasonal vaccine boosters or next-generation vaccine candidates for emerging variants.

**Fig. 4.**
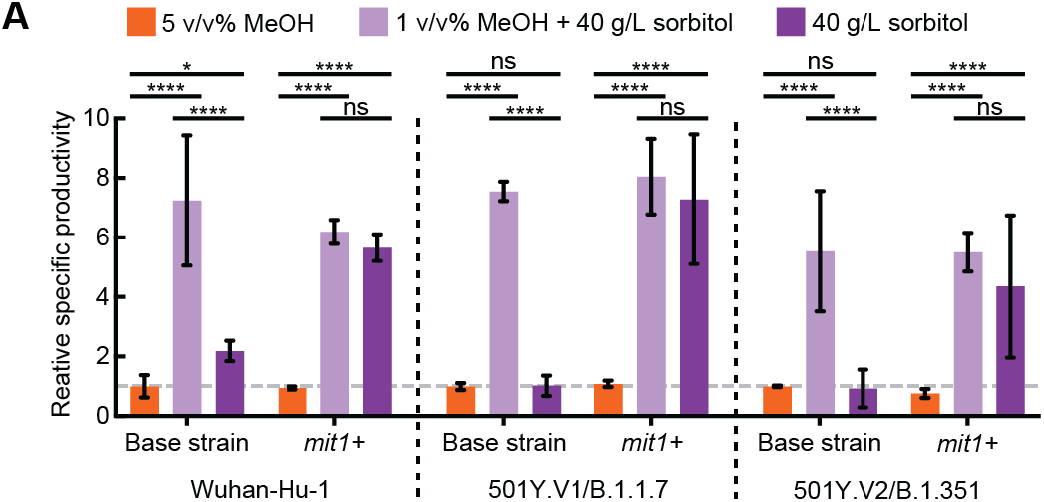
Methanol-free production of RBD variants in 3 mL culture. Error bars represent standard deviation across three biological replicates. Significance was determined by multiple t-tests with Holm Sidak correction. *p<0.01, ****p<0.000001.

In conclusion, we report here a strain that enables the manufacturing of SARS-CoV-2 RBD variants without methanol. This strain exhibits improved secreted productivity due to a reduction in protein folding stress. We demonstrated sustained productivity from the strain in a perfusion process, and scale-up to a large-scale, solvent-free fed batch process to produce a vaccine component currently in clinical trials. In this case, manufacturing at the 1,200 L scale was possible with the elimination of the requirement for methanol in the medium. Strains engineered for use without methanol and increased productivity could facilitate manufacturing of RBD and other antigens for vaccine candidates at large volumes and low costs to enable accessible and affordable vaccines for global use.

## Materials and methods

### Yeast strains

All strains were derived from wild-type *Komagataella phaffii* (NRRL Y-11430). The base strain was described previously (Brady et al., 2020). The gene containing the RBD was codon optimized, synthesized (Integrated DNA Technologies), and cloned into a custom vector. The RBD vector was transformed as described previously (Dalvie et al., 2019). Transcription factors *mit1* and *mxr1* were integrated into the genome near genomic loci GQ67_02967 and GQ67_04576, respectively, using a markerless CRISPR-Cas9 system described previously (Dalvie et al., 2019). Both *mit1* and *mxr1* were under control of the P_CAT1_ promoter from *K. phaffii*. Sequences for P_CAT1_, *mit1*, and *mxr1* were amplified from the *K. phaffii* genome.

### Cultivations

Strains for initial characterization and titer measurement were grown in 3 mL culture in 24-well deep well plates (25°C, 600 rpm), and strains for protein purification were grown in 200 mL culture in 1 L shake flasks (25°C, 250 rpm). Cells were cultivated in Rich Defined Media, described previously (Matthews et al., 2018). Cells were inoculated at 0.1 OD600, outgrown for 24 h with 4% glycerol feed, pelleted, and resuspended in fresh media with methanol or sorbitol feed to induce recombinant gene expression. Supernatant samples were collected after 24 h of production, filtered, and analyzed. InSCyT bioreactors and purification modules were operated as described previously (Crowell et al., 2018; Dalvie et al., 2021).

### Analytical assays for protein characterization

Purified protein concentrations were determined by absorbance at A280 nm. SDS-PAGE was carried out as described previously (Crowell et al., 2018). Supernatant titers were measured by reverse phase liquid chromatography as described previously (Dalvie et al., 2021), and normalized by cell density, measured by OD600. Intact mass spectrometry was performed as described previously (Dalvie et al., 2021).

### Transcriptome analysis

Cell were harvested after 18 h of production at 3 mL plate scale. RNA was extracted and purified according to the Qiagen RNeasy kit (cat #74104) and RNA quality was analyzed to ensure RNA Quality Number >6.5. RNA libraries were prepared using the 3’DGE method and sequenced on an Illumina MiSeq to generate paired reads of 20 (read 1) and 72 bp (read 2). Sequenced mRNA transcripts were demultiplexed using sample barcodes and PCR duplicates were removed by selecting one sequence read per Unique Molecular Identifier (UMI) using a custom python script. Transcripts were quantified with Salmon version 1.3.0 (Patro et al., 2017) and selective alignment using a target consisting of the *K. phaffii* transcripts, the RBD, and selectable marker transgene sequences and the *K. Phaffii* genome as a selective alignment decoy. Expression values were summarized with tximport version 1.12.3 (Soneson et al., 2016) and edgeR version 3.26.8 (McCarthy et al., 2012; Robinson et al., 2009). Expression was visualized using *log_2_(Counts per Million + 1)* values. Gene set enrichment analysis (GSEA) was performed with GSEA 4.1.0 using Wald statistics calculated by DESeq2 (Love et al., 2014) and gene sets from yeast GO Slim (Subramanian et al., 2005).

## Acknowledgements

We thank Danielle Camp for program coordination and support, Prof. Ragahavan Varadarajan of IISc for kindly providing the RBD sequence for SARS-CoV-2 Wuhan-Hu-1, and the Koch Institute’s Robert A. Swanson (1969) Biotechnology Center for technical support. This work was funded by the Bill and Melinda Gates Foundation (Investment IDs INV-002740 and INV-006131) and the AltHost Consortium. This study was also supported in part by the Koch Institute Support (core) Grant P30-CA14051 from the National Cancer Institute. The content is solely the responsibility of the authors and does not necessarily represent the official views of the NCI, the Bill and Melinda Gates Foundation, or the AltHost Consortium.

## Author Contributions

N.C.D., A.M.B., and J.C.L. conceived and planned experiments. A.M.B. and N.C.D. designed and implemented CRISPR-based genome modifications. N.C.D. and R.S.J. transformed the RBD genes. A.M.B. and C.A.W. performed RNA sequencing. A.M.B. conducted plate scale cultivations. A.M.B., S.C., and M.K.T. conducted InSCyT experiments. S.R.A. and A.M.B. performed HPLC assays. S.R.A. and L.E.C. designed and performed protein purifications. C.A.N. performed mass spectrometry. H.D.R., M.P.R., R.R.L., and U.S.S. performed and oversaw scale-up to 1,200 L fed batch. N.C.D., A.M.B., and J.C.L. wrote the manuscript. All authors reviewed the manuscript.

